# Sensitizing *Staphylococcus aureus* to antibacterial host defense by decoding and blocking the lipid flippase MprF

**DOI:** 10.1101/2020.11.12.379776

**Authors:** Christoph J. Slavetinsky, Janna N. Hauser, Cordula Gekeler, Jessica Slavetinsky, André Geyer, Alexandra Kraus, Doris Heilingbrunner, Samuel Wagner, Michael Tesar, Bernhard Krismer, Sebastian Kuhn, Christoph M. Ernst, Andreas Peschel

## Abstract

The pandemic of antibiotic resistance represents a major human health threat demanding new antimicrobial strategies. MprF is the synthase and flippase of the phospholipid lysyl-phosphatidylglycerol that increases virulence and resistance of methicillin-resistant *Staphylococcus aureus* (MRSA) and other pathogens to cationic host defense peptides and antibiotics. With the aim to design MprF inhibitors that could sensitize MRSA to both, human antimicrobials and antibiotics and support the clearance of staphylococcal infections with minimal selection pressure, we developed MprF-targeting monoclonal antibodies, which bound and blocked the MprF flippase subunit. Antibody M-C7.1 targeted a specific loop in the flippase domain that proved to be exposed at both sides of the bacterial membrane, thereby enhancing the mechanistic understanding into bacterial lipid translocation. M-C7.1 rendered MRSA susceptible to host antimicrobial peptides and antibiotics such as daptomycin. Moreover, it impaired MRSA survival in human phagocytes, which recommends MprF inhibitors for new anti-MRSA approaches. MprF-directed monoclonal antibodies provide a proof of concept for development of precisely targeted anti-virulence approaches, which block bacterial antimicrobial resistance mechanisms.

## Introduction

The continuous increase of antibiotic resistance rates undermines the significance and efficacy of available antibiotics against bacterial infections [1]. Several opportunistic antibiotic-resistant bacterial pathogens including methicillin-resistant *Staphylococcus aureus* (MRSA), vancomycin-resistant enterococci, and extended-spectrum beta-lactam or carbapenem-resistant proteobacteria impose a continuously growing pressure on modern healthcare systems [2]. MRSA is responsible for a large percentage of superficial and severe bacterial infections and the available last-resort antibiotics are much less effective than beta-lactams [3]. Unfortunately, no new class of antibiotics has entered the clinical phase since the introduction of the lipopeptide antibiotic daptomycin in 2003 [1]. Novel anti-infective strategies that would circumvent on one hand the difficulties in identifying new microbiota-preserving small-molecule antimicrobials and, on the other hand, the enormous selection pressures exerted by broad-spectrum antibiotics, are discussed as potential solutions against a looming post-antibiotic era [4]. Such strategies could be based for instance on therapeutic antibodies or bacteriophages, which usually have only a narrow activity spectrum. A possible direction could be the inhibition of bacterial targets that are of viable importance only during infection [5]. Blocking such targets by so-called anti-virulence or anti-fitness drugs would preserve microbiome integrity and create selection pressure for resistance-conferring mutations only on invading pathogens. Interfering with bacterial virulence factors should ameliorate the course of infection and enable more effective bacterial clearance by the immune system or by antibiotics.

While anti-virulence and anti-fitness therapies are discussed for several of the notoriously antibiotic-resistant bacterial pathogens, none has passed an early experimental stage, probably because most of the major pharmaceutical companies have strongly reduced their investments into new antibacterials [1, 6]. For instance, a recent approach showed promising effects reducing host colonization and invasion in meningococcal disease using phenothiazines, a family of compounds applied for treatment of psychotic disorders, to target Type IV pili, important bacterial adhesion and virulence factors [7]. Most of the documented anti-virulence approaches appear to rely on small-molecule inhibitors, which are difficult to identify and often have problematic pharmacological, toxicological, and resistance-inducing properties [6]. Monoclonal antibodies (mABs) directed against anti-virulence targets could be interesting alternatives provided the target can be reached by comparatively large antibody molecules. Therapeutic mABs are used in several malignant and inflammatory diseases [8] and have proven efficacy in toxin-mediated bacterial infections such as anthrax or clostridial toxin-mediated diseases [4, 9, 10]. Apart from toxin neutralization, however, mABs have hardly been applied in antimicrobial development programs.

The Multiple Peptide Resistance Factor (MprF), a large integral membrane protein, is crucial for the capacity of bacterial pathogens such as *S. aureus* to resist cationic antimicrobial peptides (CAMPs) of the innate immune system and CAMP-like antibiotics such as daptomycin [11–13]. MprF is highly conserved and can be found in various Gram-positive or Gram-negative pathogens [13]. MprF proteins proved to be crucial for *in vivo* virulence of various pathogens in infection models and when exposed to human phagocytes [13]. Some parts of the protein are located at the outer surface of the cytoplasmic membrane and could in principle be reached by mABs [14, 15]. MprF forms oligomers and it is a bifunctional enzyme, which can be separated into two distinct domains [15]. The C-terminal domain synthesizes positively charged lysyl-phosphatidylglycerol (LysPG) from a negatively charged phosphatidylglycerol (PG) acceptor and a Lys-tRNA donor substrate, while the N-terminal domain translocates newly synthesized LysPG from the inner to the outer leaflet of the cytoplasmic membrane [14, 16]. The exposure of LysPG at the outer surface of the membrane reduces the affinity for CAMPs and other antimicrobials [14]. Notably, *mprF* is a major hot spot for point mutations that lead to daptomycin resistance acquired during therapy of *S. aureus* infections [17].

In order to assess the suitability of MprF as a target for anti-virulence agents we developed mABs targeting several epitopes of potential extracellular loops of its transmembrane part and analyzed their capacity to bind specifically to *S. aureus* MprF. We identified a collection of mABs, which did not only bind to but also inhibited the LysPG flippase domain of MprF. Our results suggest that a specific loop between two of the transmembrane segments (TMS) of MprF is exposed at both sides of the membrane suggesting an unusual, potentially flexible topology of this protein part, which may be involved in LysPG translocation. Accordingly, targeting this loop with a specific mAB inhibited the MprF flippase function, rendered *S. aureus* susceptible to killing by antimicrobial host peptides and daptomycin, and reduced *S. aureus* survival when challenged by human CAMP-producing polymorphonuclear leukocytes (PMNs).

## Results

### 1. Generation of mABs binding to putative extracellular loops of MprF

The hydrophobic part of *S. aureus* MprF appears to include 14 TMS connected by loops with predicted lengths between two and 56 amino acids [15]. Several of the loops are located at the outer surface of the cytoplasmic membrane, accessible to mABs (Figure 1A). Peptides representing four loops with a minimum length of 13 amino acids were synthesized with N- and C-terminal cysteine residues to allow cyclization (Table S1). The peptides corresponded to three loops predicted to be at the outer membrane surface (loops one, nine, and thirteen) and loop seven, the location of which has remained ambiguous due to conflicting computational and experimental findings [15]. The N-terminal amino groups of cyclized peptides were linked to biotin to facilitate their recovery and detection. The antigen peptides were incubated with MorphoSys’s Human Combinatorial Antibody Library (HuCAL), a phage display library expressing human Fab fragments with highly diverse variable regions at the phage surface [18]. Antigen-binding phages were enriched in three iterative rounds of panning in solution and antigen-phage complexes were captured with streptavidin-coated beads. The bound phages were extensively washed to remove unspecifically binding phages, eluted, and propagated in *Escherichia coli* for a subsequent panning round. Washing steps were prolonged and antigen concentrations reduced from round one to round three to increase stringency and discard antibodies with low specificity and affinity. DNA of the eluted, antigen-specific phages was isolated and subcloned in specific *E. coli* expression vectors to yield His-tagged Fab fragments. 368 individual colonies per antigen were picked and Fab fragments were expressed and purified. A representative selection of 24 unique Fabs against all four peptides were converted to human IgG by cloning in an IgG1 expression vector system and expression in human HKB11 cells and IgGs were purified via protein A chromatography, as recently described [18] (see graphical workflow in Fig. S1).

**Figure 1.**
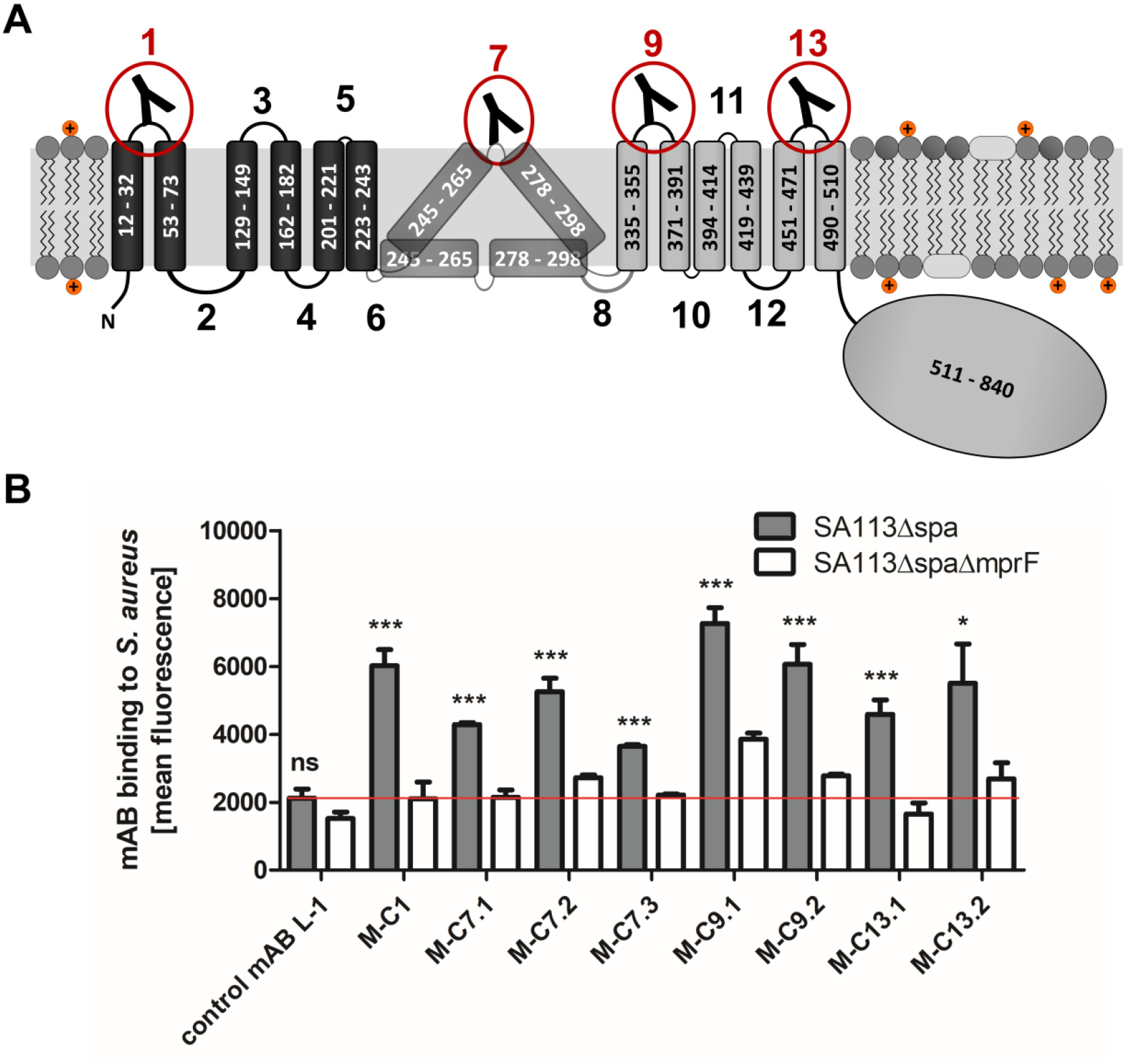
MprF topology and binding of mABs to targeted MprF-expressing *S. aureus* cells. (A) MprF membrane topology is given according to our recent study [15] showing synthase and flippase domains in gray and black, respectively. Amino acid (aa) positions of TMS and the C-terminal hydrophilic domain are indicated. TMSs from aa 245-265 and aa 278-298 are shown in two alternative positions as computational and experimental results of transmembrane topology have been contradictory [15]. Localizations of MprF’s TMS-connecting loops are numbered starting from the N-terminus, antibody-targeted loops are indicated by red circles and antibody symbols. (B) Specific binding of mABs (100 nM) to *S. aureus* was analyzed by ELISA using SA113 strains deficient in the IgG-binding protein A (Spa) comparing the SA113 *spa* mutant (*Δspa*) and *spa mprF* double knockout mutant (*ΔspaΔmprF*). The red line indicates the mean fluorescence measured at A 440 nm (affinity) of the isotype control mAB L-1 bound to *S. aureus* SA113*Δspa*. Means and SEM of at least three experiments are shown. Significant differences between SA113*Δspa* and SA113*ΔspaΔmprF* were calculated by Student’s paired t-test (ns, not significant; *, *P* < 0.05; ***, *P* < 0.0001).

The IgGs were analyzed for binding to the corresponding antigen peptides and also to the three non-cognate peptides to assess their selectivity, by ELISA with streptavidin-coated microtiter plates (Fig. S2). Antibodies directed against the four targeted MprF loops were selected based on affinity and selectivity and analyzed for binding to *S. aureus* SA113 cells expressing or not expressing MprF. Both, the “wild-type” and *mprF* deletion mutant strains lacked the gene for protein A (*spa*), which would otherwise unspecifically bind to IgG [19]. Bacteria were adsorbed to microtiter plates, blocked with bovine serum albumin (BSA), and incubated with IgGs, which were then detected with goat antihuman IgG after extensive washing. Antibodies M-C1, M-C7.1, M-C7.3, M-C9.1, M-C13.1, and M-C13.2 bound significantly stronger to MprF-expressing *S. aureus* (SA113*Δspa*) compared to MprF-deficient *S. aureus* (SA113*ΔspaΔmprF*), while the humanized isotype control mAB L-1 showed no specific binding (Fig. 1B). These findings are in agreement with the location of the loops one, nine, and thirteen at the outer surface of the cytoplasmic membrane, confirming the overall topology of MprF (Fig. 1A). Of note, loop seven between potential TMS 7 and 8 whose location had previously remained controversial was detected by antibodies M-C7.1 and M-C7.3 (Fig. 1B) indicating that loop seven is accessible from the outside. Antibodies M-C1, M-C7.1, M-C9.1, and M-C13.1 showed the strongest binding to MprF according to mean fluorescence intensities.

### 2. The MprF epitope bound by M-C7.1 is located at both, outer and inner surface of the cytoplasmic membrane

The MprF loop seven between TMS 7 and 8 bound by M-C7.1 seemed to have an ambiguous position within the cytoplasmic membrane because the flanking TMS have a comparatively low content of hydrophobic amino acids. Accordingly, only some topology analysis algorithms predict its location at the outer surface of the cytoplasmic membrane and a previous experimental topology investigation using a set of translational fusions with enzymes that are active only at intracellular or extracellular location has revealed a preferential location at the inner cytoplasmic membrane surface in *E. coli* [15]. Since our M-C7.1 and M-C7.3 binding experiments indicated accessibility of loop seven from the outside, we revisited its location in *S. aureus* with two experimental strategies.

To confirm that M-C7.1 has access to its cognate MprF antigen epitope at the outer surface of the cytoplasmic membrane in intact bacterial cells, *S. aureus* SA113*ΔspaΔmprF* expressing MprF (pRB-MprF) was grown in the presence of M-C7.1. Cells were then disrupted, membranes were solubilized by treatment with the mild non-ionic detergent n-dodecyl-β-D-maltoside (DDM), and proteins were separated in non-denaturing PAGE gels (blue native PAGE). If M-C7.1 bound before cell disruption and remained tightly attached to MprF, it should shift the MprF bands in Western blots of the blue native gels and be detectable after Western blotting of the blue native gels with a human IgG-specific secondary antibody. In order to detect MprF independently of M-C7.1 an MprF variant translationally fused to green-fluorescent protein (MprF-GFP, expressed from plasmid pRB-MprF-GFP), which is detectable by a GFP-specific primary antibody [15], was also used and treated in the same way. To detect potential non-shifted MprF proteins in the native MprF variant (pRB-MprF), M-C7.1 was used as additional primary antibody. *S. aureus* SA113*ΔspaΔmprF* bearing the empty vector (pRB) was used as a negative control. In addition to M-C7.1, all three *S. aureus* strains were also incubated with the control antibody L-1. *S. aureus* cells expressing either unmodified MprF or MprF-GFP yielded a protein band migrating at a molecular weight of around 900 kDa when preincubated with M-C7.1 but not with L-1 or in cells with the empty vector control (pRB) (Fig. 2A; Fig. S4). In contrast, the empty-vector control (pRB) strain showed an unspecific band at 300 kDa when preincubated with M-C7.1 or at 150 kDa when preincubated with L-1 but no MprF-specific band (Fig. 2A; Fig. S4). The 900-kDa band of MprF-GFP was detected by both M-C7.1 and anti-GFP confirming the identity of MprF (Fig. 2A; Fig. S4). We could recently show that MprF forms oligomers in the staphylococcal membrane, which were migrating at ca. 300 and 600 kDa [15]. Bands migrating at similar heights (ca. 250 and 500 kDa) were detected specifically in the MprF-GFP lanes after preincubation with either M-C7.1 or L-1 and only by anti-GFP primary antibodies but not by M-C7.1 as primary antibody (Fig. 2A; Fig. S4) suggesting that they represent the oligomerized but not the mAB-complexed MprF proteins [15]. Therefore, the 900-kDa bands probably represent a complex formed by MprF and M-C7.1, which confirms that M-C7.1 specifically binds MprF loop seven in live *S. aureus* cells. The fact, that M-C7.1 used as primary antibody was not able to detect 250 and 500-kDa bands of non-complexed MprF proteins neither after preincubation with M-C7.1 nor L-1 suggests that either antibody detection is prevented by the Western blot procedure or that M-C7.1 binding only occurs in the living staphylococcal cell potentially due to MprF conformational changes during flippase activity. Such conformational changes during Lys-PG translocation in live *S. aureus* would render MprF loop seven accessible to M-C7.1. Of note, the MprF-mAB complex migrating at around 900 kDa indicates that higher-order MprF multimers were shifted by complex formation with M-C7.1.

**Figure 2.**
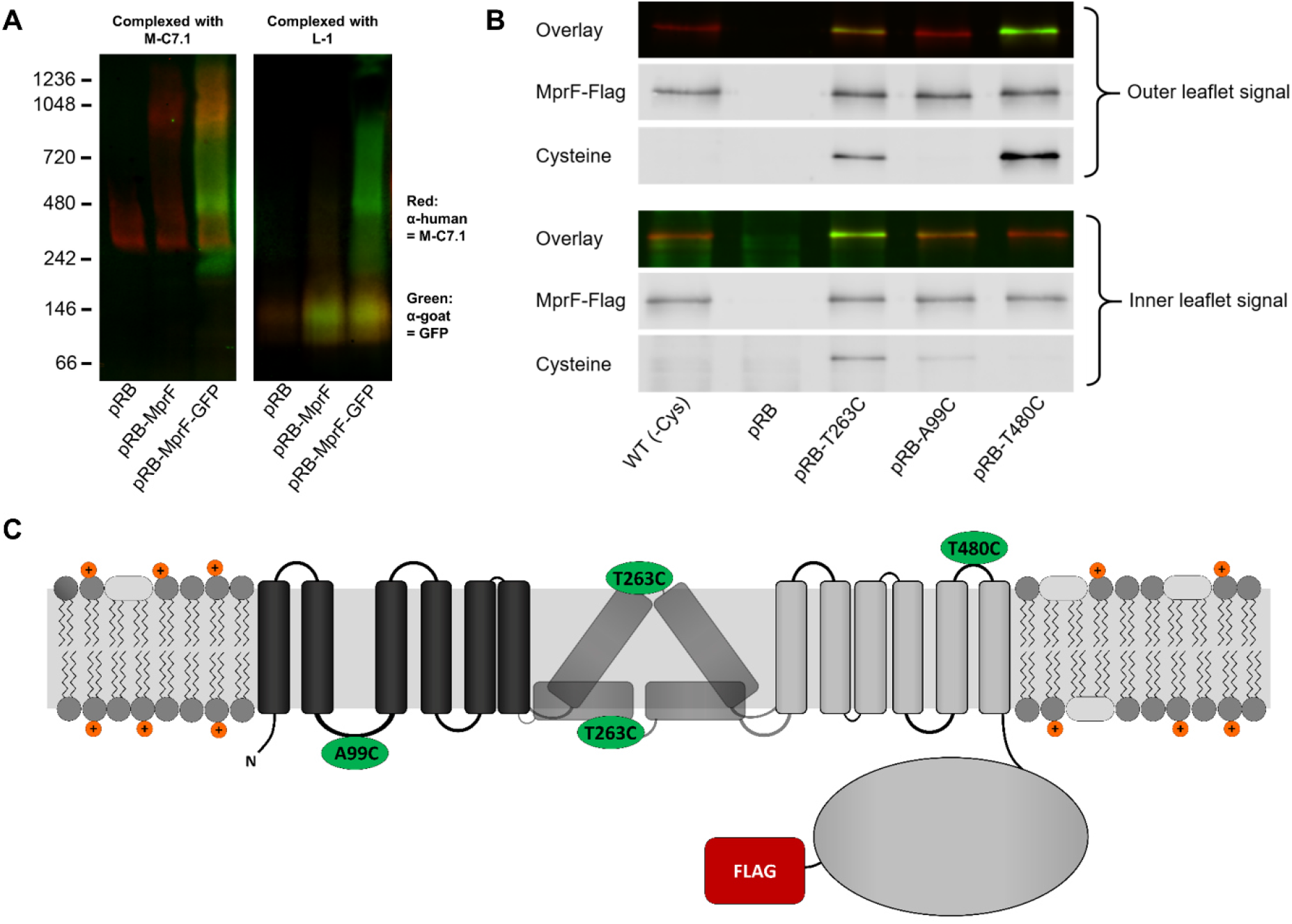
Binding of M-C7.1 to MprF and membrane localization of the M-C7.1-targeted MprF loop seven. (A) Detection of M-C7.1 binding to MprF. Plasmid-encoded native and GFP-tagged MprF variants were expressed in *S. aureus* SA113*ΔspaΔmprF* and living cells were preincubated with M-C7.1 or the isotype control mAB L-1 (in order to form MprF-mAB complexes). MprF variants/complexes were detected by blue native Western blotting via primary (anti-GFP or M-C7.1) and secondary antibodies (red for secondary antibody detection of M-C7.1; green for secondary antibody detection of anti-GFP). SA113*ΔspaΔmprF* expressing the empty vector (pRB) served as negative control. Molecular masses in kDa of marker proteins are given on the left of the gel. (B) Cellular localization of the antigen epitope of M-C7.1 using the substituted cysteine accessibility method (SCAM) for specific loops between the MprF TMSs. The substituted cysteine T263C is localized in M-C7.1’s target peptide sequence in MprF. Substitution of A99C served as inside control, substitution of T480C served as outside control (see topology model, part C). *S. aureus* SA113*ΔmprF* expressing the empty vector (pRB) and an MprF variant lacking all native cysteines (WT (-Cys)) served as additional negative controls. All MprF variants were plasmid-encoded, FLAG^®^-tagged at the C-terminus to allow immunoprecipitation and detection, and were expressed in *S. aureus* SA113*ΔmprF*. Substituted extracellular cysteine residues were labeled with Na*-(3-maleimidylpropionyl)-biocytin (MPB) (outer leaflet signal, green in overlay), while labeling of substituted internal cysteine with MPB was performed after the blocking of external cysteines with 4-acetamino-4’-maleimidylstilbene-2,2’-disulfonic acid (AMS) (inner leaflet signal, green in overlay). MprF was detected via antibody staining by an anti-FLAG^®^ antibody (red in overlay). (C) MprF topology showing location and amino acid exchanges of artificial cysteine residues for SCAM detection in green.

The position of MprF loop seven was further investigated by inserting a cysteine residue into the loop and analyzing the capacity of the membrane-impermeable agent Na*-(3-maleimidylpropionyl)-biocytin (MPB) to label cysteines covalently, using the Substituted Cysteine Accessibility Method (SCAM) [20]. MPB treatment of intact *S. aureus* cells should only lead to labelling of extracellular protein portions while blocking cysteines at the outside with 4-acetamido-4’-maleimidylstilbene-2,2’-disulfonic acid (AMS), followed by cell homogenization and addition of MPB should allow to label only protein parts at the inner surface of the membrane according to a previously established method in *E. coli* [20]. An MprF variant lacking all native cysteine residues was generated to exclude background signals. Cysteine residues in MprF were exchanged against serine or alanine residues to minimize structural or functional changes. Native *S. aureus* MprF contains six cysteines none of which is conserved in MprF proteins from other bacteria suggesting that they do not have critical functions. The cysteine-deficient mutant protein was found to be indeed functional because it decreased the susceptibility of the *S. aureus mprF* mutant (SA113*ΔmprF*) to daptomycin (Fig. S3), which depends on intact MprF synthase and flippase activities [14]. Cysteines were then inserted into MprF loop seven (T263) and, as a control, into the first intracellular loop (loop two, A99) and the last extracellular loop (loop thirteen, T480), the localization of which had been consistently confirmed by previous computational and experimental analyses [15] (Fig. 1A & 2C). Amino acids for exchange were chosen according to a predicted weak effect for functional changes by respective substitution with cysteine using https://predictprotein.org [21]. The prediction is based on a machine learning program integrating both evolutionary information and structural features such as predicted secondary structure and solvent accessibility to evaluate the effect of amino acid exchanges in a protein sequence [21]. The SCAM approach further confirmed the topology of intracellular loop two and extracellular loop thirteen (Fig. 2B), thereby demonstrating that the technique can lead to reliable results in *S. aureus*. Notably, MprF loop seven was found in both locations, at the inner and outer surface of the membrane. This finding corroborates the ambiguous position of loop seven and suggests that this loop may have some degree of mobility in the membrane, which may reflect the lipid translocation process. The finding also clarifies why M-C7.1 has the capacity to bind MprF loop seven at the outer surface of the cytoplasmic membrane.

### 3. MABs binding to putative extracellular loops of MprF render *S. aureus* susceptible to CAMPs

MprF confers resistance to cationic antimicrobials such as the bacteriocin nisin by reducing the negative net charge of the membrane outer surface [11, 14]. If the mABs would not only bind to but also inhibit the function of MprF, *S. aureus* should become more susceptible to nisin. The six mABs with confirmed specific binding to MprF and the isotype control mAB L-1 were analyzed for their capacity to increase the susceptibility of *S. aureus* SA113 to nisin. For those initial screening experiments protein A (*spa*) mutants were used to diminish effects of unspecific IgG binding to protein A. Bacteria were grown in the presence of one of the mABs and then incubated with nisin at the IC_50_ followed by quantification of viable bacterial cells. Two of the MprF-specific mABs targeting loops seven or thirteen increased the sensitivity of *S. aureus* SA113*Δspa* to nisin while the other mABs and the isotype control mAB L-1 had no significant impact (Fig. 3A). MAB M-C7.1 caused the strongest sensitization and was selected for further analysis using the highly prevalent and virulent community-associated methicillin-resistant *S. aureus* (CA-MRSA) USA300 clone [22].

**Figure 3.**
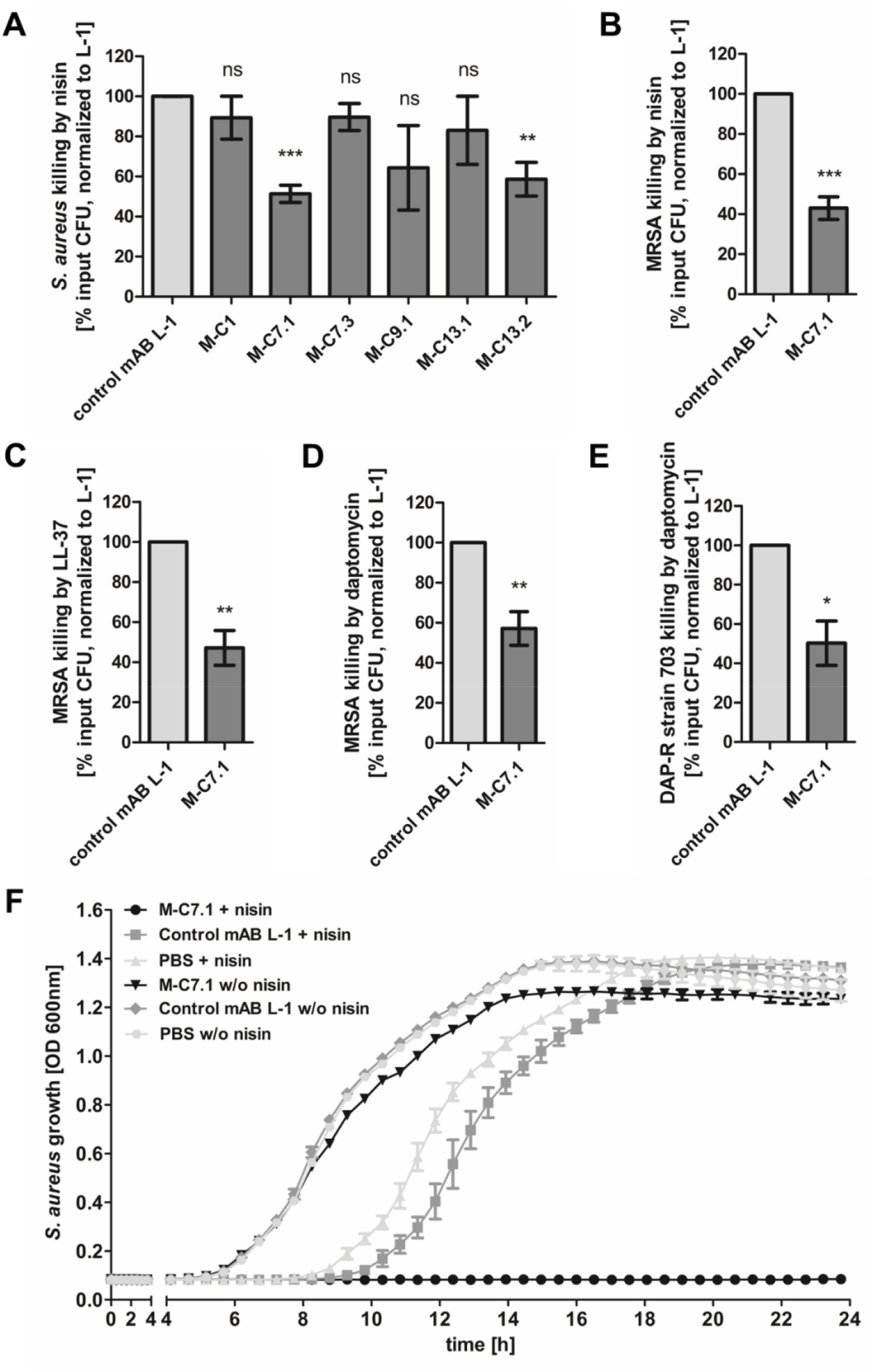
Killing and growth inhibition of *S. aureus* by antimicrobial peptides and antibiotics in the presence of M-C7.1. (A) Survival of *S. aureus* SA113*Δspa* in the presence of nisin and anti-MprF antibodies compared to control mAB L-1. Surviving CFU of each antibody (and nisin) containing sample were analyzed after 3 h incubation and the negative control (isotype mAB L-1) was set to 100% survival. (B) Survival of CA-MRSA WT strain USA300 in the presence of nisin and M-C7.1 compared to the isotype control mAB L-1. (C) Survival of USA300 WT in the presence of LL-37 and MC7.1 compared to the isotype control mAB L-1. (D) Survival of USA300 WT in the presence of daptomycin and M-C7.1 compared to the isotype control mAB L-1. (E) Survival of daptomycin-resistant CA-MRSA strain 703 [25] in the presence of daptomycin and M-C7.1 compared to the isotype control mAB L-1. (F) Growth inhibition of USA300 WT in the presence of subinhibitory nisin concentrations and M-C7.1 compared to the isotype control mAB L-1. Wells without nisin and/or antibodies served as additional negative controls. The means plus SEM of results from at least three independent experiments are shown in A-E. Technical duplicates were employed for each analysis. F shows means plus SEM of technical triplicates from a representative experiment of three independent experiments. Values for M-C7.1 or other anti-MprF antibodies that were significantly different from those for the isotype control mAB L-1, calculated by Student’s paired t-test are indicated (*, P < 0.05; **, P < 0.01; ***, P < 0.0001).

For all following experiments *S. aureus* wild type (WT) strains with intact protein A were used to make sure that the observed sensitization to CAMPs was not affected by unspecific antibody binding to protein A.

M-C7.1 was found to also increase the susceptibility of USA300 wild type to the human CAMP LL-37, a host defense peptide produced by epithelial and phagocyte cells [23], and to daptomycin, a lipopeptide antibiotic in clinical use sharing physicochemical and antibacterial properties with CAMPs [24], in addition to nisin (Fig. 3B-D). Of note, M-C7.1 also increased the antimicrobial activity of daptomycin against the daptomycin-resistant clinical MRSA isolate DAP-R 703 [25] (Fig. 3E), suggesting that M-C7.1 may potentially be able to overcome daptomycin resistance during therapy. M-C7.1 but not the isotype control mAB L-1 was able to inhibit growth of USA300 in the presence of subinhibitory concentrations of nisin (Fig. 3F). Thus, mABs specific for certain extracellular loops of MprF may not only bind to MprF but also inhibit its function.

### 4. M-C7.1 inhibits the flippase function of MprF

M-C7.1 binds MprF at the junction between the LysPG synthase and flippase domains (Fig. 1). The lipid patterns of SA113 wild type treated with M-C7.1 at concentrations that increased the susceptibility to nisin or with the isotype control mAB L-1 were compared but showed no differences indicating that the synthase function of MprF was not inhibited by M-C7.1 (Fig. 4A). The flippase activity of MprF promotes the exposure of positively charged LysPG at the outer surface of the cytoplasmic membrane thereby reducing the affinity for the small cationic protein cytochrome C [26], which binds preferentially to negatively charged PG, or for calcium-bound annexin V [27]. These model proteins have been shown to allow a sensitive assessment of changes in the surface charge of *S. aureus* in several previous studies [14, 17, 28]. Treatment of SA113 wild type with M-C7.1 at concentrations that increased the susceptibility to nisin led to a significant increase in the capacity to bind cytochrome C or annexin V compared to treatment with the isotype control mAB L-1 (Fig. 4B, C) thereby indicating that M-C7.1 inhibits the flippase function of MprF. It is tempting to speculate that the protein region of MprF loop seven and adjacent TMSs may accomplish a crucial function in the process of phospholipid translocation, as suggested by the dynamic localization of loop seven on the cytoplasmic and external faces of the membrane.

**Figure 4.**
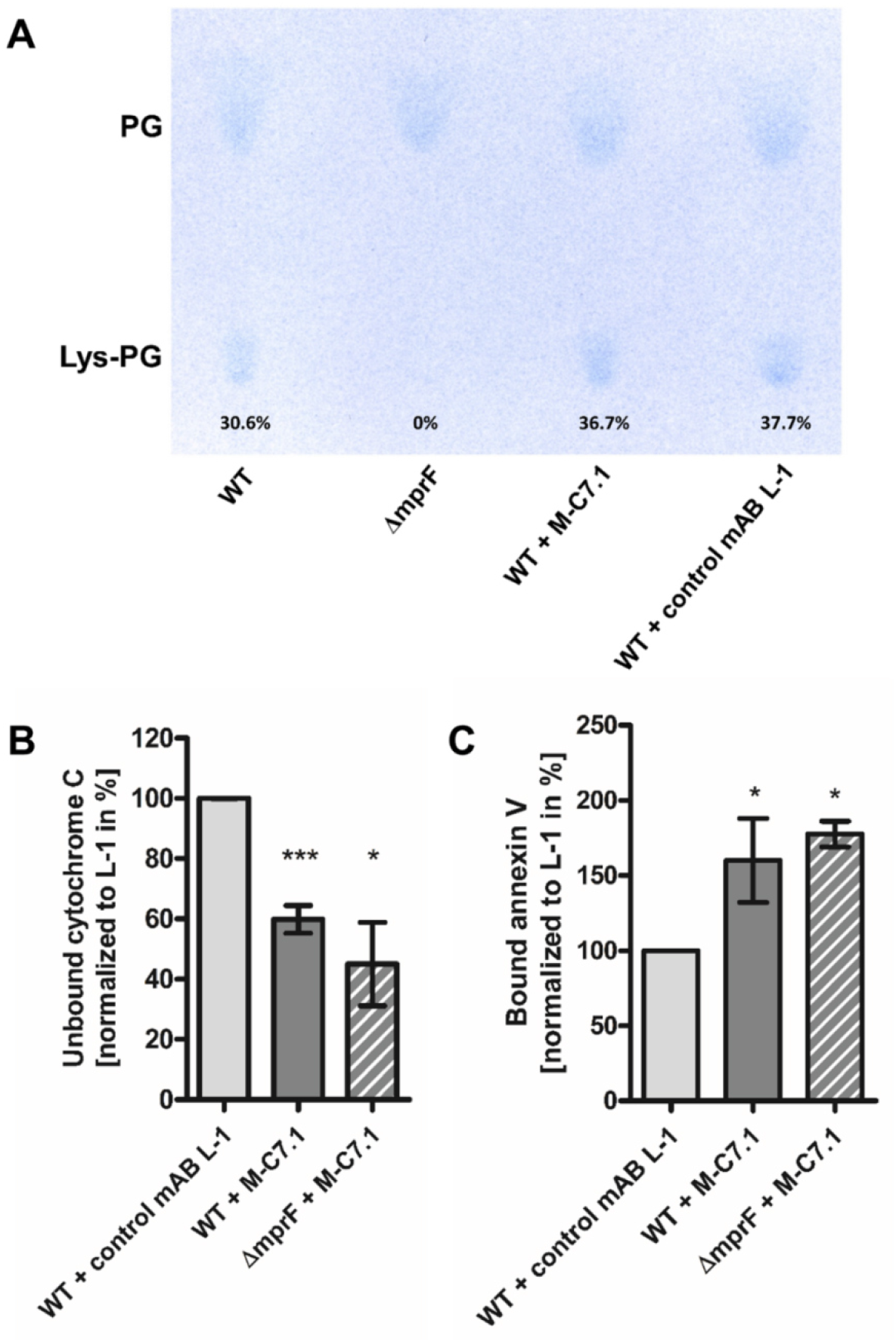
M-C7.1 inhibits the MprF LysPG flippase but not the LysPG synthase. (A) Detection of phospholipids from *S. aureus* SA113*ΔmprF* and WT treated or not treated with M-C7.1 or the isotype control mAB L-1. Polar lipids were separated by TLC and stained with the phosphate group-specific dye molybdenum blue to detect the well-documented PG and LysPG pattern of *S. aureus* WT and *mprF* deletion mutant [28]. Percentages of LysPG in relation to total phospholipid content are given below respective LysPG spots. (B) The repulsion of positively charged cytochrome C corresponds to MprF LysPG synthase plus flippase activity while the synthase activity alone does not affect repulsion. To assess MprF flippase efficiency, unbound cytochrome C in the supernatant was quantified photometrically after incubation with the *S. aureus* SA113 WT pretreated with M-C7.1 or the isotype control mAB L-1 (the latter set to 100%). SA113*ΔmprF* treated with M-C7.1 served as negative control. The means plus SEM of results from three independent experiments are shown. (C) Annexin V binding to *S. aureus* SA113 WT after incubation with M-C7.1 or the isotype control mAB L-1 was quantified by measuring cell-bound annexin V by FACS and L-1 treated samples were set to 100%. Data are expressed in relative fluorescence units. The means plus SEM of results from three independent experiments are shown in B and C. Significant differences between M-C7.1 and L-1-treated samples were calculated by Student’s paired t-test (*, P < 0.05; ***, P < 0.0001).

### 5. M-C7.1-treatment abrogates *S. aureus* survival in phagocytes

The capacity of PMNs to kill phagocytosed bacteria does not only rely on the oxidative burst but also on the activity of LL-37 and other CAMPs and antimicrobial proteins [29]. The increased susceptibility of M-C7.1-treated *S. aureus* to CAMPs should therefore alter its survival ability in PMNs. CA-MRSA wild-type strain USA300 was pretreated with mABs, opsonized with normal human serum, and exposed to human PMNs. Treatment with M-C7.1 or the isotype control mAB L-1 did not alter the rate of PMN phagocytosis, but M-C7.1-treated USA300 cells were significantly more rapidly killed by PMNs than those treated with the isotype control antibody (Fig. 5A, B). This finding reflects our previous reports on reduced survival of MprF-deficient *S. aureus* in PMNs [11, 30]. Thus, *S. aureus* treatment with M-C7.1 might reduce the capacity to persist in infections and may help to blunt the virulence of *S. aureus* in invasive infections.

**Figure 5.**
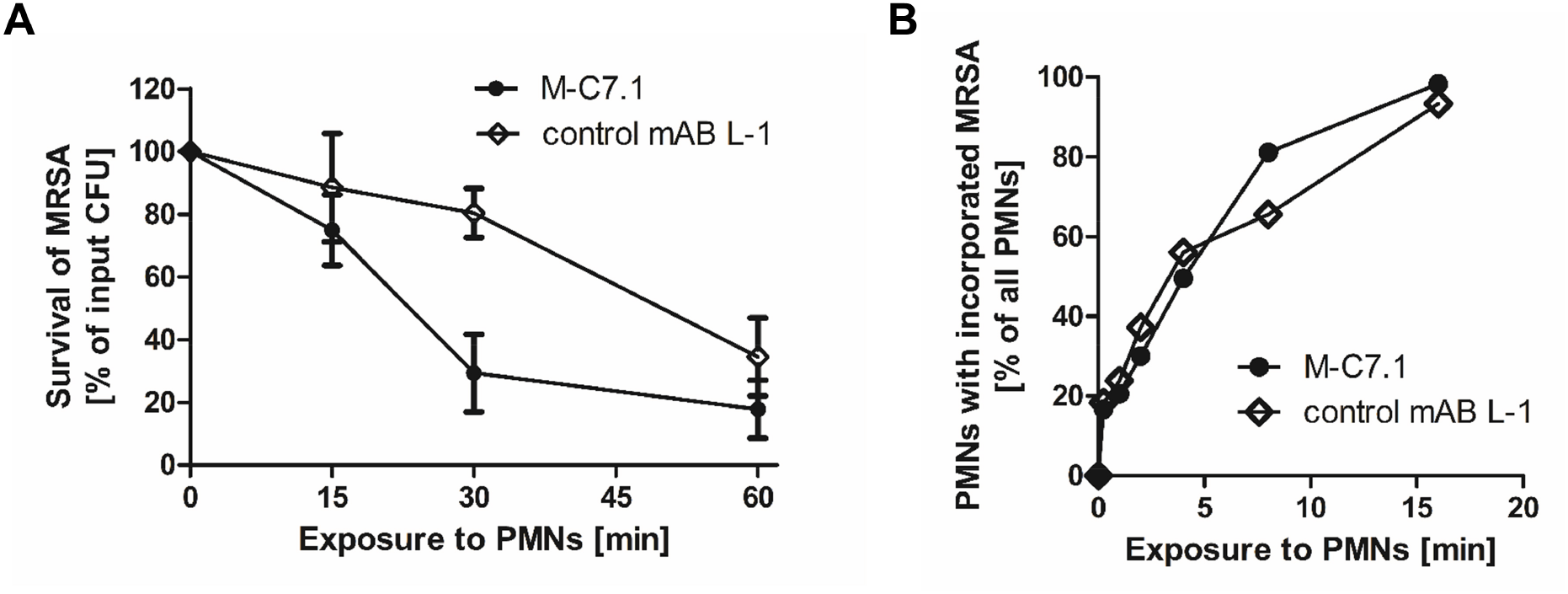
M-C7.1 supports *S. aureus* clearance by isolated human PMNs. (A) Kinetics of killing of CA-MRSA strain USA300 WT treated with M-C7.1 compared to isotype control mAB L-1 by freshly isolated human PMNs. Viable bacteria (CFU) after incubation with PMNs are shown as percentage of initial CFU counts. The means plus SEM of results from three independent experiments are shown. (B) Kinetics of phagocytosis of USA300 WT treated with M-C7.1 compared to isotype control mAB L-1 by freshly isolated human PMNs. Percentages of PMNs bearing FITC-labeled USA300 are given. Means of three counts from a representative experiment are shown.

## Discussion

Monoclonal therapeutic antibodies have proven efficacy for neutralization of bacterial toxins such as *Clostridium botulinum* or *Clostridioides difficile* toxins, and mAB-based therapies targeting the *S. aureus* alpha toxins and leukotoxins are currently developed [4, 31]. Moreover, therapeutic mABs binding to *S. aureus* surface molecules to promote opsonic phagocytosis have been assessed in preclinical and clinical trials [32]. In contrast, mABs inhibiting crucial cellular mechanisms of bacterial pathogens have hardly been assessed so far [33]. We developed specific mABs, which can block the activity of *S. aureus* MprF, the first described bacterial phospholipid flippase. Our mABs did not mediate increased internalization of *S. aureus* cells, most probably because the bacterial cell wall is too thick to allow binding of the mAB FC part to phagocyte FC receptors. However, the inhibition of MprF by M-C7.1 sensitized *S. aureus* to CAMPs and daptomycin and promoted killing by human PMNs, which use CAMPs as an important component of their antimicrobial arsenal. Specific inhibition of MprF could therefore promote the capacity of human host defense to clear or prevent a *S. aureus* infection and would, at the same time, increase the susceptibility of *S. aureus* to CAMP-like antibiotics such as daptomycin. Thus, targeting bacterial defense mechanisms provides a promising concept for anti-virulence therapy. M-C7.1 sensitized S. aureus also in the presence of protein A indicating that the unspecific IgG-binding by protein A does not interfere with the M-C7.1 capacity to block MprF.

Specific binding of MprF antibody M-C7.1 to the extracellular loop seven inhibited the flippase function of MprF, which indicates that this loop plays a crucial role in the lipid translocation process. Loop seven is located between predicted TMS 7 and 8, and its presence at the outer surface of the cytoplasmic membrane had remained elusive since computational and experimental analyses had yielded conflicting results [15]. Surprisingly, we found loop seven to be accessible from both, the outside and inside of the cytoplasmic membrane. This suggests that the protein part formed by loop seven and adjacent TMS 7 and 8 may change its position in the protein complex, moving between the two membrane surfaces to accomplish LysPG translocation (Fig. 6), or it could be in a unique position of the large MprF protein complex that allows access from both sides. This protein part might represent the center of the MprF flippase establishing the previously reported MprF-dependent distribution of charged LysPG in the inner and outer leaflet of the cytoplasmic membrane of *S. aureus* [14] (Fig. 6). Interestingly, in living staphylococcal cells, M-C7.1 bound to putative higher-order MprF multimers as shown via blue native Western blotting suggesting that such multimers might represent the active lipid translocating state of MprF. Replacement of conserved amino acids in the M-C7.1-targeted protein part has been shown to inhibit the flippase function of MprF [15], which underscores the crucial function of this domain. Notably, one of the conserved and essential amino acids in TMS 7, D254, is negatively charged. It is tempting to speculate that D254 may interact with the positively charged head group of LysPG during the translocation process. Mutations on loop seven and adjacent TMS have recently been found to cause specific resistance to the structurally related lipopeptide antibiotics daptomycin and friulimicin B but not to other antimicrobials [17]. Indirect evidence has suggested that these mutations allow the flippase to translocate these lipopeptide antibiotics instead of or together with LysPG [17], which is in agreement with a direct role of loop seven in the translocation process. A mAB binding loop thirteen also sensitized *S. aureus* to nisin in a similar but less pronounced way as M-C7.1 suggesting that this loop may also have a critical role for MprF activity.

**Figure 6.**
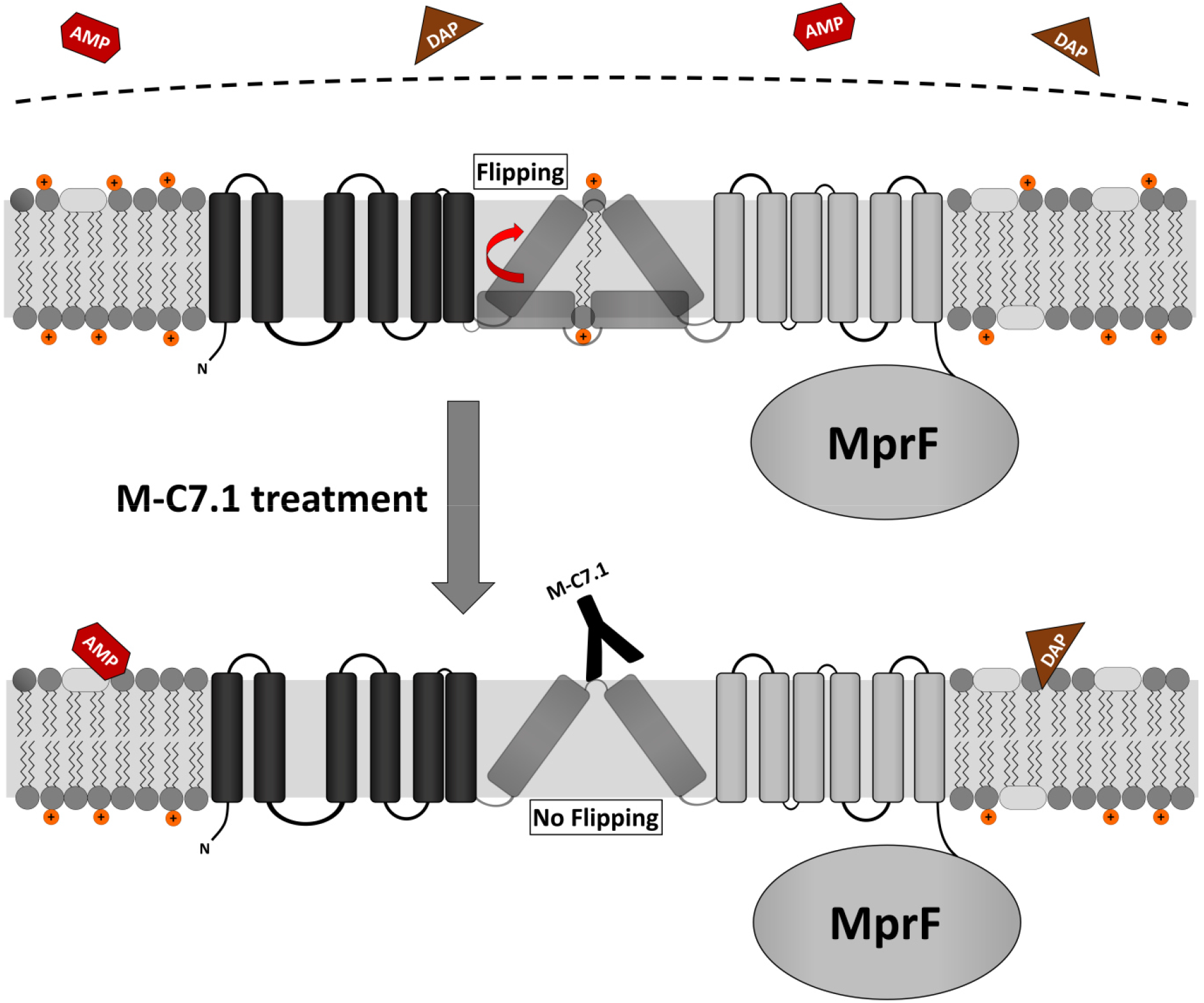
Proposed model for MprF inhibition by M-C7.1. Flipping of positively charged LysPG probably by TMS 7 and 8 of MprF results in a more positively charged staphylococcal membrane able to repulse AMPs or daptomycin (DAP). Binding of M-C7.1 to MprF loop seven blocks the flippase, which results in a more negatively charged staphylococcal membrane and subsequently in an increased *S. aureus* membrane disruption by AMPs and daptomycin.

With the severe shortage of new antibiotic target and drug candidates, crucial cellular machineries conferring fitness benefits during infection rather than accomplishing essential cellular functions should be considered as targets for the development of new therapeutics. The phospholipid synthase and flippase MprF may become a role model for further targeting of bacterial defense mechanisms for such anti-fitness or anti-virulence drugs. Compared to other potential targets, MprF has the advantage that essential parts of it are exposed at the external surface of the cytoplasmic membrane. Moreover, MprF is found in many other bacterial pathogens for which similar therapeutic mABs could be developed [14]. On the other hand, only a few amino acids in loop seven are conserved in other bacteria, which would limit the impact of loop-seven-directed mABs on other microbiome members thereby minimizing the resistance selection pressure and the risk of dysbiosis.

## Materials and Methods

### Bacterial strains, maintenance and mutagenesis of mprF

We used commonly used strains, the methicillin-susceptible laboratory strain *S. aureus* SA113 (ATCC 35556), methicillin-resistant clinical clone *S. aureus* USA300, and the SA113 *mprF* knock-out derivative SA113*ΔmprF*, which has been described recently [26, 34] (Tab. S2).

For the construction of a protein A mutant (Δ*spa*) the *E. coli/S. aureus* shuttle vector pKOR1 [35] was used, which allows allelic replacement with inducible counter-selection in staphylococci. Flanking regions of *spa* were amplified from chromosomal DNA of *S. aureus* COL with primer pairs Spa-del_attB1 (ggggacaagtttgtacaaaaaagcaggc**caatattccatggtccagaact; bold: spa sequence**) and Spa-del for BglII (gtcg*agatct*ataaaaacaaacaatacacaacg, restriction site italic) as well as Spa-del_attB2 (ggggaccactttgtacaagaaagctggg**atcagcaagaaaacacacttcc; bold: spa flanking sequence**) and Spa-del rev BglII (aaa*agatct*aacgaattatgtattgcaata, restriction site italic). Both PCR products were digested with BglII and subsequently ligated. Without further purification, the ligation product was mixed with equimolar amounts of pKOR1 and *in vitro* recombination was performed with BP clonase Mix (Invitrogen) according to the manufacturer’s instructions. The recombination mixture was transferred to chemically competent *E. coli* DH5α and isolated plasmids from the resulting transformants were analyzed by restriction digest. The correct plasmid was isolated from *E. coli* DH5α and used to electroporate competent *S. aureus* RN4220 from which it was again isolated and transferred into *S. aureus* SA113 by electroporation. Allelic replacement was essentially conducted as previously described [35] and resulting deletion mutants were confirmed by PCR.

The *S. aureus* SA113 *spa mprF* double mutant (SA113*ΔspaΔmprF*) was constructed by transducing the gene deletion cassette of the *S. aureus* SA113 *mprF* deletion mutant (SA113*ΔmprF*) to the marker-less *spa*-deficient *S. aureus* SA113 mutant (SA113*Δspa*) using standard transduction protocols. The resulting *S. aureus* strain *SA113ΔspaΔmprF* was identified by screening for erythromycin resistance conferred by the gene deletion cassette and confirmed by PCR of the deleted genome section. Bacteria were maintained on tryptic soy agar plates.

Cysteine substitutions in *mprF* were accomplished by site-directed mutagenesis in *Escherichia coli* using *E. coli/S. aureus* shuttle vector pRB474 bearing *mprF*, using the QuickChange II Site-Directed Mutagenesis Kit (Agilent), as described recently [17]. Mutated *mprF* derivatives in pRB474*mprF* were transferred into SA113*ΔmprF* and *mprF* expression was mediated by the plasmid-encoded constitutive *Bacillus subtilis* promoter *vegII*. 10 μg/ml chloramphenicol served for maintenance in all plasmid-based studies. A99, T263, and T480 were chosen for exchange against a cysteine residue because of prediction of a weak effect on protein structure in an analysis of functional changes given a single point mutation according to https://predictprotein.org [21].

Plasmids and primers used in this study are given in Table S3 and S4, respectively. Unless otherwise stated, bacteria were cultivated in Mueller-Hinton Broth (MHB, Sigma-Aldrich) with appropriate antibiotics for all experiments.

### Antigen selection and antibody production

MprF-derived peptides of interest (Fig. 1 and Tab. S1) were custom-synthesized by JPT Peptide Technologies GmbH (Berlin). An N-terminal and a C-terminal cysteine were added to enable cyclization. Biotin was coupled to the peptide via a 4,7,10-trioxa-1,13-tridecanediamine succinic acid (Ttds)-linker [36]. Cyclization and coupling of linker and biotin were performed by JPT Peptide Technologies.

Recombinant antibodies were generated from the HuCAL PLATINUM^®^ library, as described recently, by three iterative rounds of panning on the antigen peptides in solution and antigen-antibody-phage complexes were captured with streptavidin-coated beads (Dynal M-280) [18, 37]. Fab-encoding inserts of the selected HuCAL PLATINUM phagemids were cloned and expressed in *E. coli* TG1 F-cells, purified chromatographically by IMAC (Bio-Rad) and subcloned in an IgG1 expression vector system to obtain full-length IgG1s by expression in eukaryotic HKB11 cells, as described recently [18]. IgGs were purified by protein A affinity chromatography (MabSelect SURE, GE Healthcare) [18].

The humanized antibody MOR03207 [38] directed against chicken lysozyme serves as isotype control and was called L-1 in this manuscript.

### Peptide and whole-cell ELISAs of MprF specific mABs

Overnight *S. aureus* SA113*Δspa* or SA113*ΔspaΔmprF* cultures grown in MHB were adjusted to OD_600_ 0.1 in fresh MHB and grown to OD_600_ of at least 1.0. Cells were harvested, adjusted to OD_600_ 0.5 with 0.9% NaCl, and used to coat 96 well NUNC Maxisorp Immuno Plates for 1 hour (50 μl/well). After three washing steps with Tris-buffered saline (TBS; 50 mM Tris, 150 mM NaCl in 800 mL of H_2_O, pH 7.4) containing 0.05% Tween 20 (TBS-T), the cells were blocked with PBS containing 1 x ROTI^®^Block (Carl Roth) for 1 h. Primary antibodies were added after three washing steps with TBS-T at an indicated final concentration of 1, 10 or 100 nM and incubated for 1 h. After three more washing steps with TBS-T, anti-human IgG conjugated to alkaline phosphatase (Sigma-Aldrich A8542) diluted 1:10,000 in TBS-T was applied. Finally, after three washing steps with 2x TBS-T and 1x TBS, the cells were incubated with 100 μl p-nitrophenyl phosphate solution as advised by the manufacturer (Sigma-Aldrich N1891). Fluorescence was measured at an excitation of 440±25 nm and an emission of 550±35 nm.

### Detection of antibody binding to MprF by blue native Western blotting

To detect binding of the anti-MprF antibody to MprF we performed blue native polyacrylamide gel electrophoresis (BN-PAGE) as previously described [15] with slight modifications. Briefly, 20 ml of *S. aureus* cells grown in MHB with anti-MprF antibodies (100 μg/ml) to OD_600_ 1 were incubated with 750 μl of lysis buffer (100 mM EDTA [pH 8.0], 1 mM MgCl_2_, 5 μg/ml lysostaphin, 10 μg/ml DNase, proteinase inhibitor [cocktail set III from Calbiochem] diluted 1:100 in PBS) for 30 min at 37°C and homogenized three times with 500 μl zirconia-silica beads (0.1 mm diameter from Carl-Roth) using a FastPrep 24 homogenizer (MP Biomedicals) for 30 s at a speed of 6.5 m/s. After removing the beads, cell lysate was centrifuged 20 min at 14,000 rpm and 4°C to sediment cell debris and supernatant was transferred to microcentrifuge polypropylene tubes (Beckman Coulter) and cytoplasmic membranes were precipitated by ultracentrifugation for 45 min at 55,000 rpm and 4°C (Beckman Coulter rotor TLA 55). Membrane fractions resuspended in resuspension buffer (750 mM aminocaproic acid, 50 mM Bis-Tris [pH 7.0]) were incubated with the MprF- or lysozyme-specific humanized IgG mAB and/or a GFP-specific rabbit IgG mAB (Invitrogen) for 30 min shaking at 37°C. Dodecyl maltoside was added to a final concentration of 1% for 1 h at 4°C to solubilize MprF-antibody complexes. Insoluble material was removed by ultracentrifugation for 30 min at 40,000 rpm and 4°C. 20 μl supernatant was mixed 1:10 with 10 × BN loading dye (5% [wt/vol] Serva Blue G, 250 mM aminocaproic acid, 50% glycerol) and run on a Novex NativePAGE 4 to 16% Bis-Tris gel (Invitrogen). Separated proteins were transferred to a polyvinyl difluoride membrane and detected via either, goat anti-human IgG DyLight 700 (Pierce) or goat anti-rabbit IgG DyLight 800 (Pierce) as secondary antibodies in an Odyssey imaging system from LI-COR.

### Substituted cysteine accessibility method (SCAM) to localize MprF loops in inner and/or outer membrane leaflet

SCAM was adapted for use in *S. aureus* from the protocol recently established for *E. coli* [20]. Variants of *S. aureus* SA113*ΔmprF* were expressing cysteine-deficient and/or -altered MprF derivatives bearing a FLAG^®^ tag at the C-terminus via the plasmid pRB474. Bacteria were grown at 37°C to an OD_600_ 0.7-0.8 in 100 ml TSB, split into two equal aliquots, and harvested. Pellets were resuspended in a mixture of buffer A (100 mM HEPES, 250 mM sucrose, 25 mM MgCl_2_, 0.1 mM KCl [pH 7.5]) supplemented with 1 mM MgSO_4_, 1 mM EDTA, 2.1 μg lysostaphin, 0.7 μg DNase and 1% of a protease inhibitor cocktail (Roche) and incubated for 45 min at 37°C. For staining of cysteine residues in the outer leaflet of the cytoplasmic membrane, cells of the first aliquot were treated with N*-(3-maleimidylpropionyl) biocytin (MPB, Thermo Scientific) at a final concentration of 107 μM for 20 min on ice. In the second aliquot, external cysteine residues were blocked with 107 μM 4-acetamino-4’-maleimidylstilbene-2,2’-disulfonic acid (AMS, Thermo Scientific) for 20 min on ice, bacterial cells were subsequently disrupted in a bead mill as described above and internal cysteines were labeled with MPB for 5 min. MPB labeling was quenched in both aliquots by the addition of 21 mM β-mercaptoethanol. Cell debris was removed by several centrifugation and washing steps, membrane fractions were collected via ultracentrifugation at 38,000 x g, and labeled membrane samples were stored at −80°C.

For protein enrichment and precipitation, labeled membrane samples were thawed on ice and suspended in 20 mM β-mercaptoethanol containing buffer A. Proteins were solubilized by the addition of an equal volume of solubilization buffer (50 mM Tris-HCl [pH 9], 1 mM EDTA, 2% SDS) and vigorous vortexing for 30 min at 4°C, followed by an incubation step of 30 min at 37°C and vortexing for 30 min at 4°C. After solubilization, one and a half volumes of immunoprecipitation buffer 1 (50 mM Tris-HCl [pH 9], 150 mM NaCl, 1 mM EDTA, 2% Thesit, 0.4 % SDS [pH 8.1]) was added and samples were incubated for 2.5 h with magnetic FLAG^®^ beads (Sigma-Aldrich) in a rotation wheel at 4°C. After several washing steps with immunoprecipitation buffer 1 and immunoprecipitation buffer 2 (50 mM Tris-HCl [pH 9], 1 M NaCl, 1 mM EDTA, 2% Thesit, 0.4 % SDS [pH 8.1]), FLAG^®^-tagged MprF was eluted by the addition of 0.1 M glycine-HCl (pH 2.2) and neutralized by adding Tris-HCl (pH 9).

For detection of cysteine-labelled MprF, samples were analyzed by denaturing SDS-PAGE and Western blotting. To this end, 12-μl samples were mixed with 4 x Laemmli sample buffer (Bio-Rad), loaded onto a 10%-polyacrylamide gel, and separated via electrophoresis. Proteins were transferred onto a polyvinylidene difluoride (PVDF) membrane (Immobilon-PSQ PVDF membrane, Merck) by semi-dry turbo Western blot procedure (Trans-Blot Turbo Transfer System, Bio-Rad). FLAG^®^-tagged MprF was detected both by a mouse anti-FLAG^®^ primary antibody (Sigma-Aldrich) and goat antimouse secondary antibody (LI-COR) at 700 nm, while MBP-labelled cysteine residues were detected with streptavidin DyLight conjugate (Thermo Scientific) at 800 nm in an Odyssey imaging system from LI-COR.

### Determination of susceptibility to antimicrobial agents

Overnight cultures of *S. aureus* SA113*Δspa* or USA300 WT were diluted in fresh MHB and adjusted to OD_600_ 0.25 (~1.5 x 10^7^ cells). Antibodies were adjusted to a concentration of 1 mg/ml and 10 μl per well of a 96 well plate were added to 90 μl of the adjusted cell suspension (final antibody concentration: 100 μg/ml). Cells were grown in the presence of the isotype control antibody L-1 or with the respective anti-MprF antibodies. After 3 hours of incubation at 37°C under shaking, optical density was determined, and cells were adjusted to OD_600_ 0.025 in 500 μl cold PBS. 80 μl of the adjusted cell suspension were mixed with 20 μl of antimicrobial substances to final concentrations of 22.5 μg/ml nisin for SA113*Δspa* or of 5 μg/ml nisin for USA300, 1.5 μg/ml daptomycin for USA300, 11 μg/ml daptomycin for daptomycin resistant CA-MRSA strain 703, and 45 μg/ml LL-37 for USA300 or 20 μl PBS as control. After incubation for 2 hours under shaking at 37°C, the cell suspensions were diluted 1:2,000 and 100 μl of each duplicate was plated in triplicates on TSB agar plates to obtain a representative value of bacterial survival.

MICs of daptomycin were determined using MIC test strips (Liofilchem) according to the manufacturer’s advice.

To analyze the capacity of mAB M-C7.1 to inhibit *S. aureus* growth, overnight cultures of CA-MRSA strain USA300 grown in MHB medium were diluted to OD_600_ 0.01 with fresh MHB medium containing 1 μM of antibodies and 4 μg/ml nisin (Sigma-Aldrich) or PBS as a control in 96 well plates. Plates were incubated for 24 hours at 37°C under shaking conditions using a Bioscreen C (Oy Growth Curves Ab Ltd). Optical density at 600 nm was determined and compared to growth without nisin.

### Isolation and quantification of polar lipids

*S. aureus* cultures (1 ml in MHB) were grown to the exponential phase (OD_600_ 1) in the presence of antibodies (final concentration: 100 μg/ml) as described above for the determination of susceptibility to antimicrobials. Polar lipids were extracted using the Bligh-Dyer method [39], with chloroform/methanol/sodium acetate buffer (20 mM) (1:1:1, by vol.), vacuum-dried, and resuspended in chloroform/methanol (2:1, by vol.). Extracts were filled into a 100-μl Hamilton syringe and spotted onto silica gel 60 F254 high-performance thin-layer chromatography (HPTLC) plates (Merck) with a Linomat 5 sample application unit (Camag) and run in a developing chamber ADC 2 (Camag) with a running solution composed of chloroform-methanol-water (65:25:4, by vol.). Phosphate group-containing lipids were detected by molybdenum blue staining and phospholipids were quantified in relation to total phospholipid content by determining lipid spot intensities densitometrically with ImageJ (http://rsbweb.nih.gov/ij/docs/guide/index.html) as described recently [28].

### Determination of S. aureus cell surface charge

For analysis of the cytochrome C repulsion capacity of *S. aureus*, bacterial cells were grown in the presence of antibodies (100 μg/ml) as described above for the analysis of antimicrobial susceptibility. Bacteria were adjusted to OD_600_ 1 in 1 ml PBS, pelleted, resuspended in 100 μl 0.05 mg/ml cytochrome C (Sigma-Aldrich) followed by incubation at 37°C under shaking. The cells were then pelleted by centrifugation and absorption of the supernatant containing unbound cytochrome C was determined at OD_410_

To determine annexin V binding to *S. aureus*, bacterial cells were grown in the presence of antibodies (100 μg/ml) as described above at 37°C with shaking for 3 hours, harvested, washed twice in PBS buffer, and resuspended in PBS containing CaCl_2_ to OD_600_ 0.5 in 1 ml. Bacteria were gently mixed with 5 μl of allophycocyanin (APC)-labeled annexin V (Thermo Scientific) and incubated at room temperature for 15 min in the dark, as described recently [17]. At least 50,000 bacterial cells per antibody were analyzed by flow cytometry to quantify surface-bound fluorophore (FL-4).

### Phagocytosis and killing of S. aureus by human polymorphonuclear leukocytes (PMNs)

Human PMNs are the major human phagocytes to counteract bacterial infection and form the largest subgroup (~95%) of PMNs [29]. PMNs were isolated from fresh human blood of healthy volunteers by standard Ficoll/Histopaque gradient centrifugation as described recently [40]. Cells were resuspended in HBSS containing 0.05% human serum albumin (HBSS-HSA; HBSS with 0.05% Albiomin^®^, Biotest AG).

CA-MRSA strain USA300 was prepared for PMN experiments as described for the determination of susceptibility to antimicrobials, grown in the presence of antibodies (100 μg/ml) at 37°C under shaking for 3 hours, harvested, washed twice in HBSS, and adjusted to a density of 5 × 10^7^ CFU/ml. Pooled serum from healthy human volunteers (blood bank of the University Hospital Tübingen) was added to a final concentration of 5% and bacteria were opsonized for 10 min at 37°C, as described recently [11]. Prewarmed bacteria and PMNs were mixed to final concentrations of 5 × 10^6^ CFU/ml and 2.5 × 10^6^ PMNs/ml in flat-bottom 96 well plates together with antibodies (final concentration 100 μg/ml). Samples of 50 μl were shaken at 37°C.

For analysis of the PMN killing capacity, incubation was stopped at different time points by the addition of 100-fold volumes of ice-cold distilled water to disrupt the PMNs. Triplets of appropriate sample volumes were spread on LB agar plates and colonies were counted after 24 h incubation at 37°C. Numbers of live bacteria did not change during the 60-min incubation period in the absence of PMNs compared to the initial bacteria counts.

For phagocytosis studies, overnight cultures of bacteria were labeled with 0.1 mg/ml fluorescein-5-isothiocyanate (FITC) at 37°C for 1 h, as described previously [11]. After washing with PBS, the bacteria were resuspended in HBSS-HSA, adjusted to 5 × 10^6^ CFU/ml, mixed with PMNs (2.5 × 10^6^ PMNs/ml), and opsonized as described above (shaking at 37°C). Incubation was stopped by addition of 100 μl ice-cold 1% paraformaldehyde. The percentage of PMNs bearing FITC-labeled bacteria was determined by flow cytometric analysis of 20,000 cells.

### Statistics

Statistical analyses were performed with the Prism 5.0 package (GraphPad Software) and Group differences were analyzed for significance with the two-tailed Student’s *t* test. A *P* value of ≤0.05 was considered statistically significant.

## Author contributions

C.J.S, A.K., S.W., M.T., B.K., C.M.E., and A.P. conceived and designed the experiments. C.J.S., J.N.H., C.G., J.S., A.G., D.H., B.K., and C.M.E. performed the experiments. C.J.S., J.N.H., J.S., A.G., A.K., S.W., S.K., C.M.E., and A.P. analyzed the data. S.W., M.T., and A.P. contributed material / analysis tools. C.J.S., J.N.H., S.W., C.M.E., and A.P. wrote the paper.

## Acknowledgements

This work was financed by Grants from the German Research Foundation TRR34, SFB766, TRR156, TRR261, and GRK1708 to A.P.; and the German Center of Infection Research (DZIF) to A.P. C.J.S. was supported by the intramural Experimental Medicine program of the Medical Faculty at the University of Tübingen and received grants from the DZIF and the cluster of Excellence EXC2124 Controlling Microbes to Fight Infection (CMFI). The authors acknowledge infrastructural support by the cluster of Excellence EXC2124 CMFI.

## Supplemental figures

**Supplemental Figure S1.**
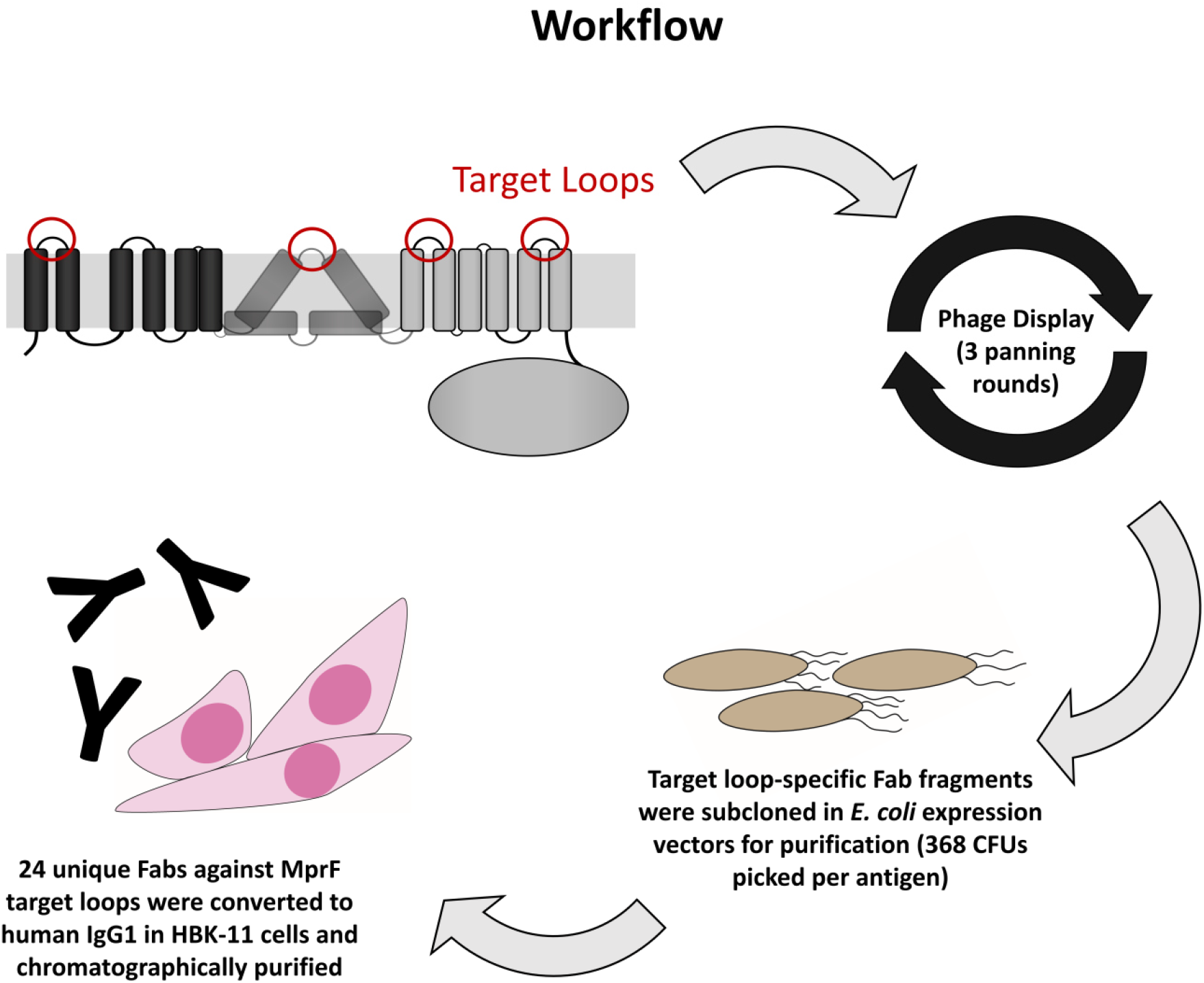
Workflow for the development of MprF specific mABs. Biotinylated MprF peptide loops were incubated with the HuCal phage display library expressing single-chain human Fab fragments [18], antigen-binding phages were enriched in three iterative panning rounds, bound antigen-specific phages were isolated and respective Fab fragments were subcloned in *E. coli* expression vectors to yield His-tagged Fab fragments, 24 unique Fabs against all peptides were converted to human IgG by cloning in an IgG1 expression vector system for human HKB11 cells, and IgGs were purified via protein A chromatography.

**Supplemental Figure S2.**
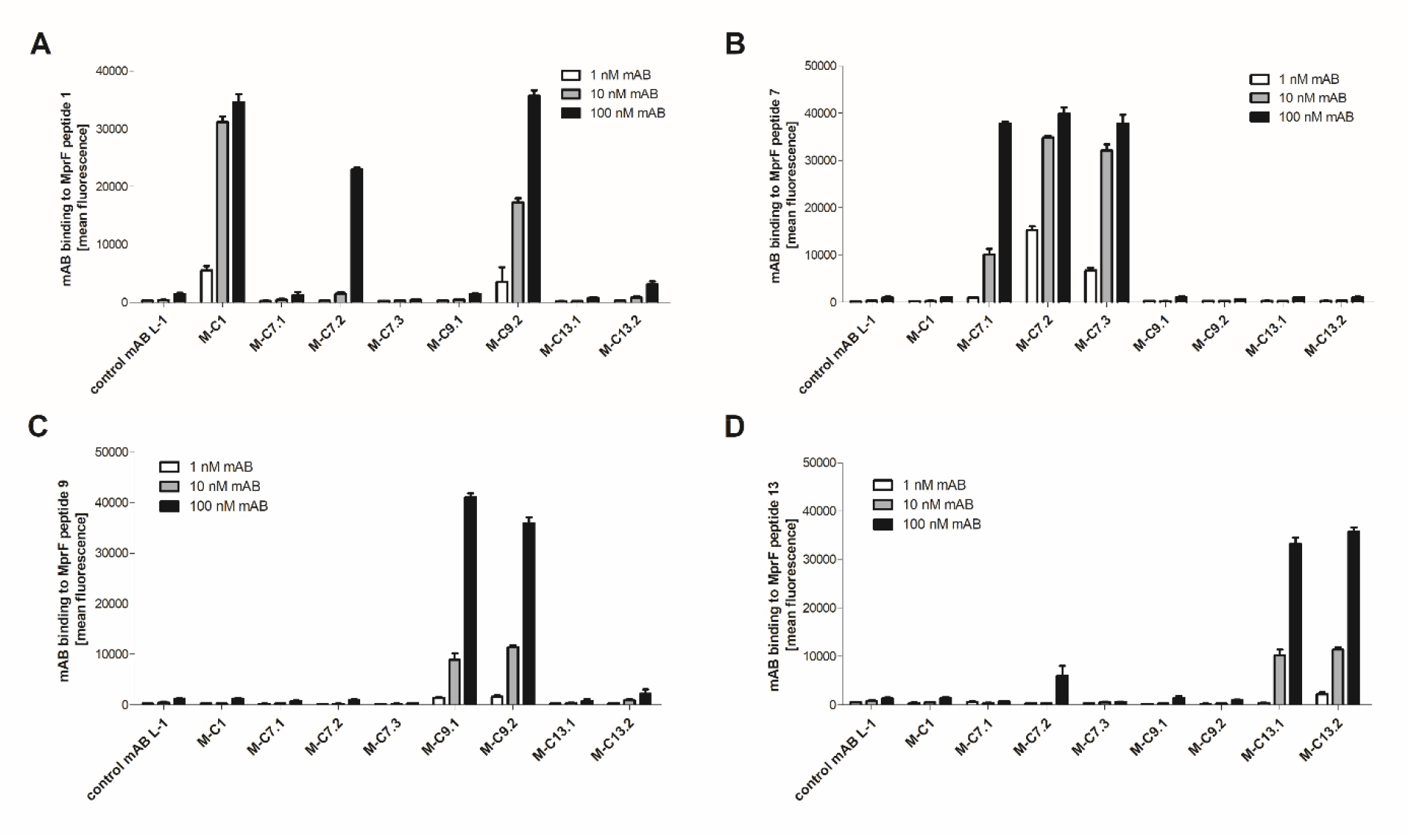
Specific binding of selected mABs to cyclic MprF-derived target peptides was analyzed by ELISA. Biotinylated cyclic peptides corresponding to the MprF loops 1, 7, 9, and 13 were incubated with eight anti-MprF IgGs in PBS compared to the control mAB L-1. (A-D) show binding of mABs at increasing concentrations to cyclic MprF loops 1, 7, 9, and 13, respectively. Mean fluorescence measured at A 440 nm and SEM of three independent experiments are shown.

**Supplemental Figure S3.**
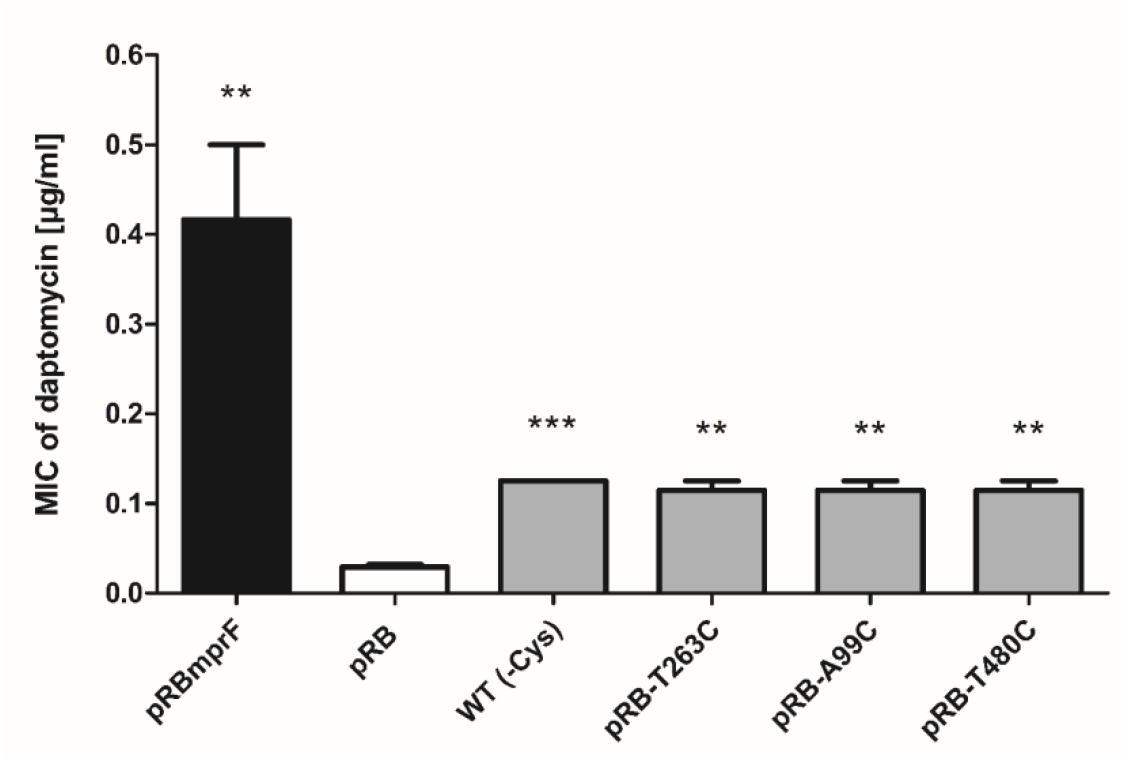
Effects of cysteine replacement and insertion on MprF function, assessed by measuring daptomycin susceptibility. Minimal inhibitory concentrations (MICs) of daptomycin against the indicated *S. aureus* strains are shown. The *mprF* deletion mutant with empty pRB474 plasmid served as a negative control. The means plus SEM of results from three independent experiments are shown. Values that are significantly different from the values determined for *S. aureus* SA113*ΔmprF* bearing pRB474 (pRB), calculated by Student’s paired t-test, are indicated (**, *P*<0.01; ***, *P*< 0.0001).

**Supplemental Figure S4.**
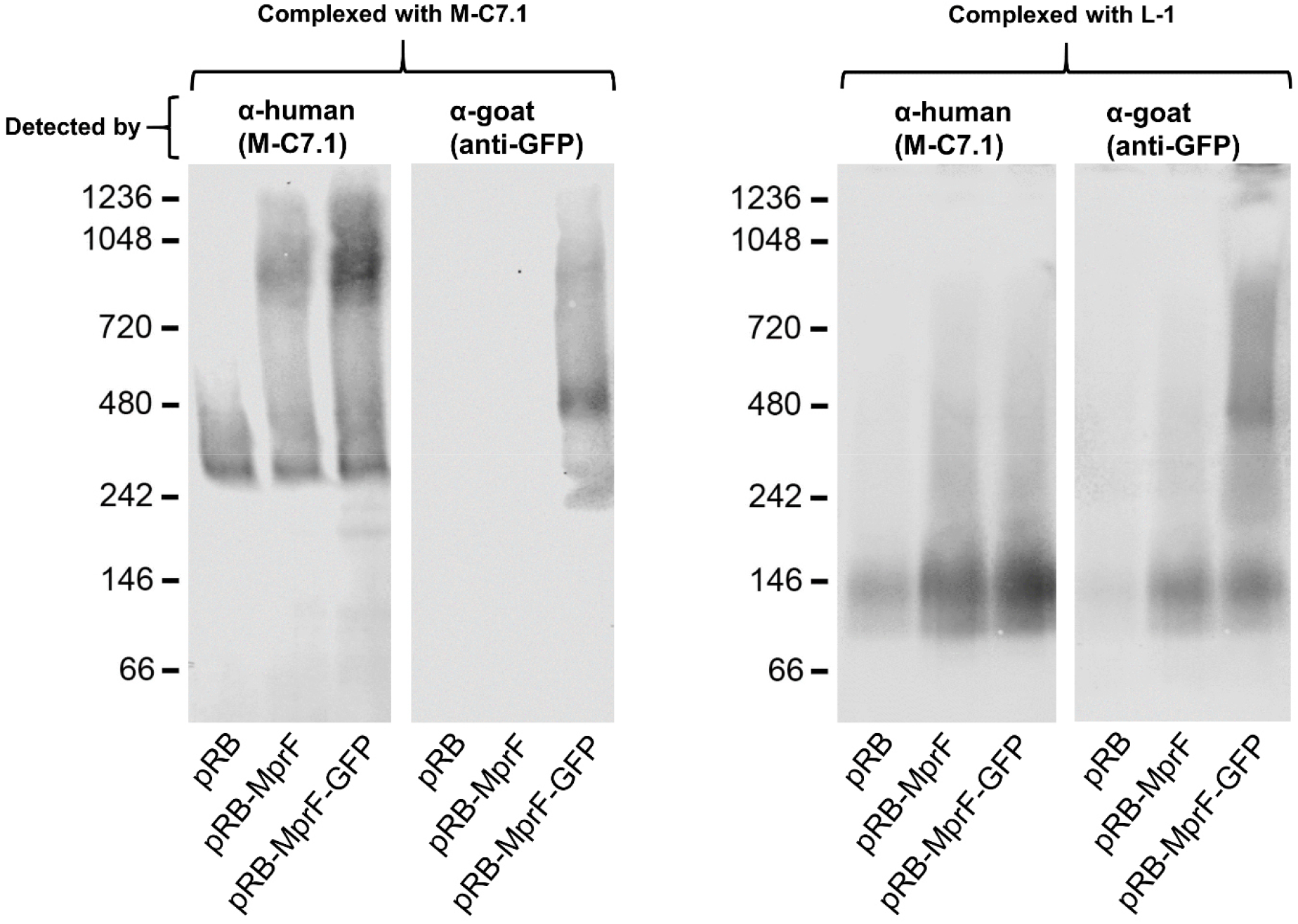
Detection of M-C7.1 binding to MprF. Single channels are showed in black and white from figure 3A. Further explanations are found in figure legend of figure 3A.

## Supplemental tables

**Table S1.**
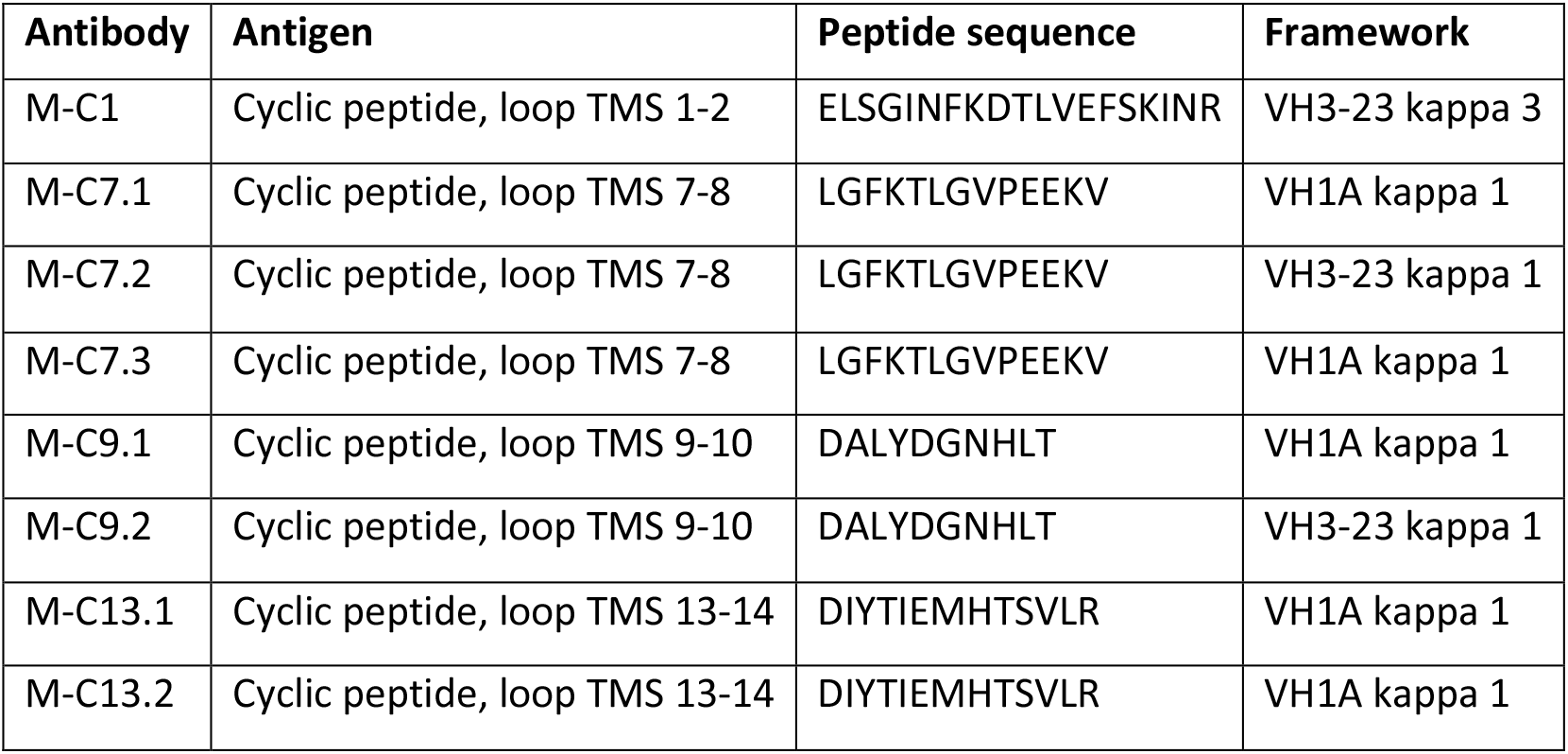
MprF-directed antibodies and its target peptides.

| Antibody | Antigen | Peptide sequence | Framework |
| --- | --- | --- | --- |
| M-C1 | Cyclic peptide, loop TMS 1-2 | ELSGINFKDTLVEFSKINR | VH3-23 kappa 3 |
| M-C7.1 | Cyclic peptide, loop TMS 7-8 | LGFKTLGVPEEKV | VH1A kappa 1 |
| M-C7.2 | Cyclic peptide, loop TMS 7-8 | LGFKTLGVPEEKV | VH3-23 kappa 1 |
| M-C7.3 | Cyclic peptide, loop TMS 7-8 | LGFKTLGVPEEKV | VH1A kappa 1 |
| M-C9.1 | Cyclic peptide, loop TMS 9-10 | DALYDGNHLT | VH1A kappa 1 |
| M-C9.2 | Cyclic peptide, loop TMS 9-10 | DALYDGNHLT | VH3-23 kappa 1 |
| M-C13.1 | Cyclic peptide, loop TMS 13-14 | DIYTIEMHTSVLR | VH1A kappa 1 |
| M-C13.2 | Cyclic peptide, loop TMS 13-14 | DIYTIEMHTSVLR | VH1A kappa 1 |

**Table S2.** Bacterial strains used in this study.

| Strain name | Characteristics |
| --- | --- |
| *S. aureus* SA113 WT (ATCC 35556) | Restriction-deficient *S. aureus* strain derived from NCTC 8325 [41] |
| *S. aureus* SA113Δ*mprF* | MprF deletion mutant of SA113, Erm<sup>r</sup> [11] |
| *S. aureus* SA113Δ*spa* | Spa deletion mutant of SA113. Constructed in this study. Markerless. |
| *S. aureus* SA113Δ*spa*Δ*mprF* | Spa and MprF double deletion mutant of SA113. Constructed in this study. Erm<sup>r</sup> |
| *S. aureus* DAP-R MRSA 703 | Daptomycin resistant clinical MRSA isolate possessing a single point mutation in *mprF* (S295L) [25] |
| *S. aureus* USA300 LAC | CA-MRSA wild type strain [34] |
| *E. coli* TG1 | Strain for phage display usage [18] |

**Table S3.**
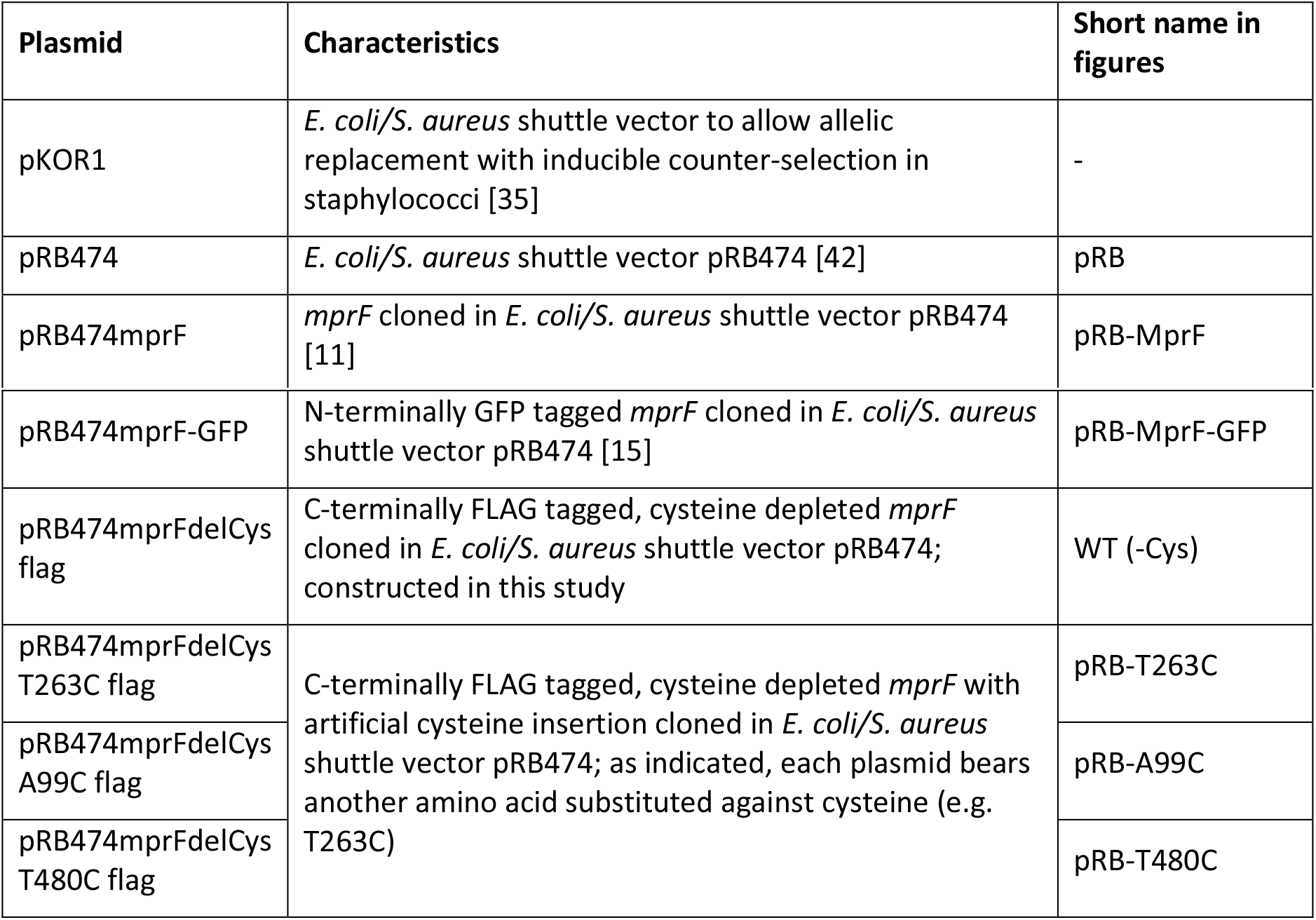
Plasmids used in this study.

| Plasmid | Characteristics | Short name in figures |
| --- | --- | --- |
| pKOR1 | *E. coli*/*S. aureus* shuttle vector to allow allelic replacement with inducible counter-selection in staphylococci [35] | - |
| pRB474 | *E. coli*/*S. aureus* shuttle vector pRB474 [42] | pRB |
| pRB474*mprF* | *mprF* cloned in *E. coli*/*S. aureus* shuttle vector pRB474 [11] | pRB-MprF |
| pRB474mprF-GFP | N-terminally GFP tagged *mprF* cloned in *E. coli/S. aureus* shuttle vector pRB474 [15] | pRB-MprF-GFP |
| pRB474mprFdelCys flag | C-terminally FLAG tagged, cysteine depleted *mprF* cloned in *E. coli/S. aureus* shuttle vector pRB474; constructed in this study | WT (-Cys) |
| pRB474mprFdelCys T263C flag | C-terminally FLAG tagged, cysteine depleted *mprF* with artificial cysteine insertion cloned in *E. coli/S. aureus* shuttle vector pRB474; as indicated, each plasmid bears another amino acid substituted against cysteine (e.g. T263C) | pRB-T263C |
| pRB474mprFdelCys A99C flag |  | pRB-A99C |
| pRB474mprFdelCys T480C flag |  | pRB-T480C |

**Table S4.**
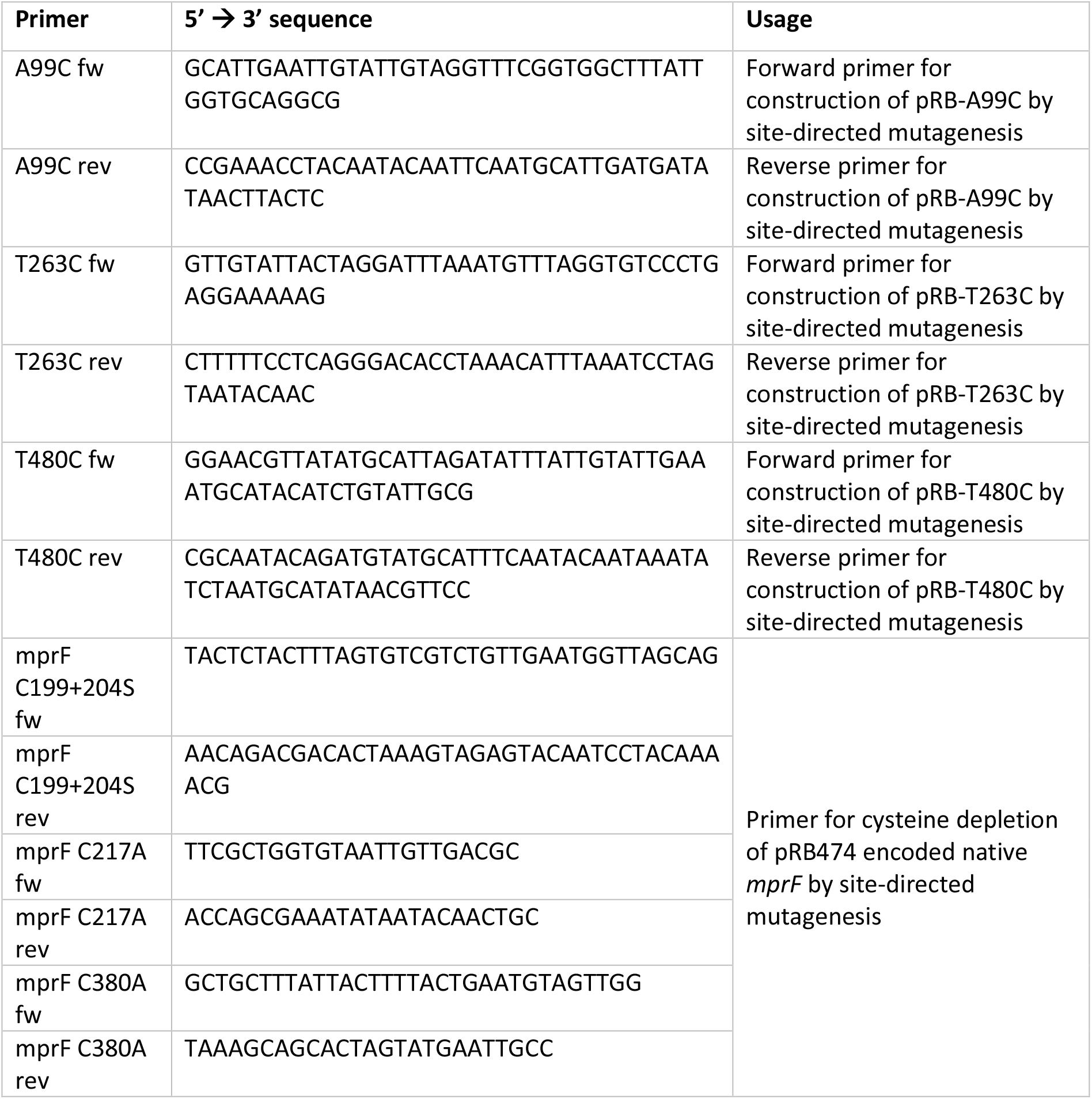
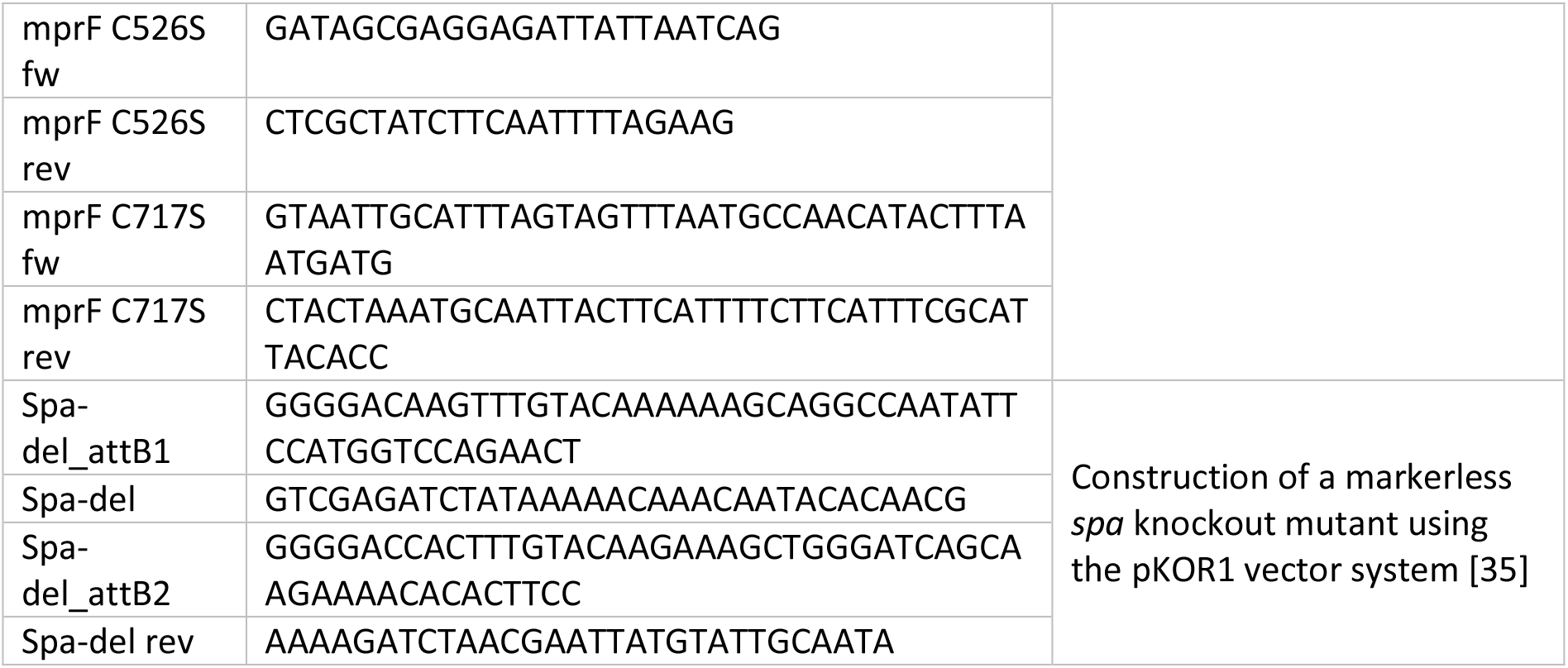
Primers used in this study.

